# Ultra-sensitive mutation detection and genome-wide DNA copy number reconstruction by error corrected circulating tumour DNA sequencing

**DOI:** 10.1101/213306

**Authors:** Sonia Mansukhani, Louise J. Barber, Sing Yu Moorcraft, Michael Davidson, Andrew Woolston, Beatrice Griffiths, Kerry Fenwick, Bram Herman, Nik Matthews, Ben O’Leary, Sanna Hulkki, David Gonzalez De Castro, Michael Hubank, Anisha Patel, Andrew Wotherspoon, Aleruchi Okachi, Isma Rana, Ruwaida Begum, Matthew Davies, Thomas Powles, Katharina von Loga, Nick Turner, David Watkins, Ian Chau, David Cunningham, Naureen Starling, Marco Gerlinger

## Abstract

Minimally invasive circulating free DNA (cfDNA) analysis can portray cancer genome landscapes but highly sensitive and specific genetic approaches are necessary to accurately detect mutations with often low variant frequencies. We developed a targeted cfDNA sequencing technology using novel off-the-shelf molecular barcodes for error correction, in combination with custom solution hybrid capture enrichment. Modelling based on cfDNA yields from 58 patients shows that our assay, which requires 25ng of cfDNA input, should be applicable to >95% of patients with metastatic colorectal cancer. Sequencing of a 163.3 kb target region including 32 genes detected 100% of single nucleotide variants with 0.15% variant frequency in cfDNA spike-in experiments. Molecular barcode error correction reduced false positive mutation calls by 98.6%. In a series of 28 patients with metastatic colorectal cancers, 80 out of 91 (88%) mutations previously detected by tumour tissue sequencing were called in the cfDNA. Call rates were similar for single nucleotide variants and small insertions/deletions. Mutations only called in cfDNA but not detectable in matched tumour tissue included, among others, a subclonal resistance driver mutation to anti-EGFR antibodies in the *KRAS* gene, multiple activating *PIK3CA* mutations in each of two patients (indicative of parallel evolution), and *TP53* mutations originating from clonal haematopoiesis. Furthermore, we demonstrate that cfDNA off-target read analysis allows the reconstruction of genome wide copy number aberration profiles from 71% of these 28 cases. This error-corrected ultra-deep cfDNA sequencing assay with a target region that can be readily customized enables broad insights into cancer genomes and evolution.

## Introduction

Many cancers release cell free DNA (cfDNA) into the circulating blood and numerous publications have demonstrated the feasibility to detect tumour somatic mutations, copy number changes and genomic rearrangements from this cfDNA (1–6). Such liquid biopsies can provide critical insights into tumour genomic landscapes, for example to inform precision medicine approaches such as tailored therapeutic interventions (7), or to predict recurrences after surgery (8, 9). The minimally invasive nature of cfDNA analysis also enables longitudinal tracking of cancer genomic changes without the discomfort, costs and risk associated with invasive tumour biopsies. Finally, sampling of cancer genomes from the blood provides the opportunity to not only detect clonal but also subclonal mutations that are often missed by biopsies as a consequence of spatial intratumour heterogeneity and the resultant sampling biases (10, 11).

Although there are clear advantages of tumour genomic analyses from cfDNA, the low tumour-derived cfDNA fraction found in many cancers, and the even lower abundances of subclonal mutations, complicate the detection of somatic mutations in plasma. Genetic techniques with high sensitivity and low false positive error rates are hence crucial for reliable cfDNA analysis. Digital droplet PCR (ddPCR) and BEAMing assays can accurately detect point mutations present at frequencies below 0.1% in cfDNA but are restricted to the analysis of a limited number of genomic loci (12, 13). This precludes their use to survey larger genomic regions. In contrast, next generation sequencing (NGS) can interrogate larger regions such as the entire coding region of genes included in target gene panels. However, NGS error rates rapidly increase when attempting to call mutations with variant allele frequencies (VAFs) below 5% (14). Error correction approaches have been incorporated into NGS cfDNA assays to reduce this error rate, for example through background error correction models that remove recurrent false positives occurring at specific positions and more recently by including random unique molecular identifiers/barcodes (MBC) combined with redundant over-sequencing to remove PCR amplification and sequencing errors (15, 16). The latter approach has been shown to enable accurate mutation calling with VAFs ≤0.1%. However, these methods use proprietary rather than off-the-shelf reagents and this limits their application to answer research questions requiring the design of customized cfDNA sequencing panels.

In the presented study, we assessed how novel, commercially available, MBC reagents combined with customized solution hybrid capture target enrichment technology can be optimized for ultra-deep error-corrected cfDNA sequencing. We describe the optimisation and the technical performance of this cfDNA sequencing technology, including the ability to detect VAFs close to 0.1%. We subsequently assess the concordance of mutation calls from cfDNA with those called by clinical grade tumour tissue sequencing in patients with metastatic colorectal cancer (mCRC). We demonstrate a high sensitivity (88%) to call tumour mutations through cfDNA sequencing and that additional driver mutations, likely resulting from ongoing cancer evolution, can be detected. Finally, we show that genome-wide copy number profiles can be simultaneously inferred from cfDNA by analysing off-target reads.

## Results

### Determining the optimal ctDNA quantity for metastatic colorectal cancer analysis

cfDNA was extracted from the plasma of 58 patients with metastatic colorectal cancer (mCRC) enrolled in the FOrMAT trial at the Royal Marsden Hospital (ClinicalTrials.gov NCT02112357) (17). Plasma samples had either been collected from treatment naïve patients (n=19) or at the time tumours started to progress after prior palliative systemic treatment (n=39). cfDNA is naturally fragmented with a predominant fragment size of approximately 160 bp (18). DNA fragments with multiples of this size can also be found but these are less abundant. We quantified cfDNA with a size range from 100 to 700 bp to encompass these fragment peaks (Figure 1A). A median of 20.8 ng cfDNA per ml of plasma were extracted (25^th^ percentile: 7.1 ng per ml; 75^th^ percentile: 49.1 ng per ml) with a minimum of 1.03 ng per ml (Figure 1B).

**Figure 1.**
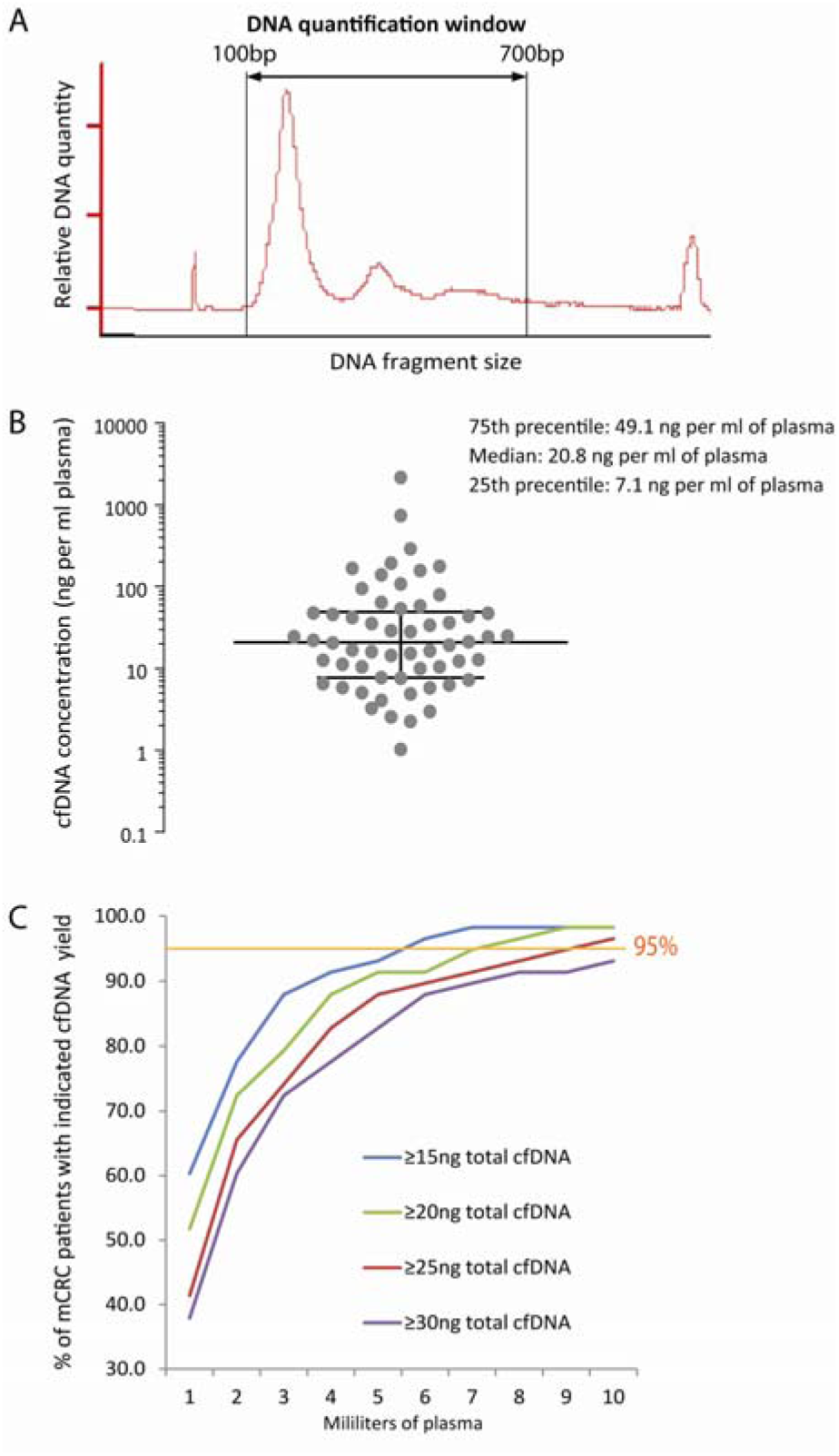
(A) Example of a cfDNA sample Bioanalyzer profile. cfDNA was quantified across a window from 100 bp to 700 bp in all samples in this study. (B) cfDNA yields (ng per ml plasma) obtained from 58 patients with metastatic colorectal cancer. (C) Fraction of patients that achieve the indicated total cfDNA yield (see legend) based on the plasma volume available from each patient.

We next wanted to define an optimized input cfDNA quantity that can be obtained reliably from a standard blood draw of 20-30 ml from at least 95% of mCRC patients. We used the cfDNA yield data to model how the plasma volume that is available per patient influences the fraction of patients from which at least 15, 20, 25 or 30 ng of total cfDNA can be obtained (Figure 1C). This showed that 25 ng of cfDNA can be obtained from over 95% of mCRC patients if 10 ml of plasma are collected. This plasma volume can be reliably obtained from a blood draw of 20-30 ml. Furthermore, with a haploid human genome mass of approximately 3.3 pg, 25 ng of cfDNA should contain in excess of 7500 haploid genome equivalents. This is theoretically sufficient for the detection of mutations with frequencies around 0.1% VAF, even if only 20% of the cfDNA fragments are incorporated into sequencing libraries as expected from studies with off-the-shelf solution hybrid capture library preparation reagents (19). Based on these results, we chose 25 ng as the standard input quantity for our cfDNA assay.

### Optimization of sequencing library preparation and of cfDNA sequencing

We designed a custom solution hybrid capture panel targeting 32 genes including CRC driver genes and other genes of interest for on-going studies, with a total target region of 163.3 kb (details in Supplementary Table 1). The Agilent XT HS library preparation kit, which tags every individual DNA strand with a random 10 base molecular barcode (MBC), was used for sequencing library preparation. Due to the short fragment size of cfDNA molecules, no additional fragmentation was used. Pre- and post-capture PCRs were optimized so that, from 25 ng standard input cfDNA, they reliably provided sufficient yields for target-DNA capture (>500 ng) and for sequencing (>200 nM) with a minimal number of PCR cycles (see methods section for details of the optimized protocol).

Libraries were prepared from 7 patient and 2 healthy donor cfDNA samples with the manufacturer’s original library preparation protocol that recommended a post-capture wash temperature of 65 °C. These were sequenced in 75 bp paired-end rapid output mode on an Illumina HiSeq2500 sequencer with a median of 80,453,088 reads per sample. The on target fraction varied between samples, with only a median of 30% of reads aligned to the target region across this small series. This showed a low efficacy of target capture in many cases, possibly as a consequence of using a small target panel or low input DNA quantities (20). Therefore, we assessed whether the on-target fraction could be increased by using an alternative protocol from the manufacturer, which washed the DNA bound to capture beads after hybridisation more stringently with a temperature of 70 °C instead of 65 °C. To test this, two library preparations were started in parallel from each of four cfDNA samples and the post-capture wash was performed at 65 °C in one library and at 70 °C in the other. After sequencing with similar read numbers (65°C: median of 92,820,887 per sample; 70°C: median of 102,582,694 per sample) the on-target fraction significantly (p<0.01) increased to 71-74% with the 70°C protocol compared to 30-35% with 65°C (Figure 2A). Down-sampling of the 70°C data to match the total reads generated for the 65°C samples did not affect the on-target fraction (data not shown). Hence, the more stringent 70°C wash reduced the non-specific carry over of DNA fragments into the final library and this optimized library preparation method was chosen as our standard protocol.

**Figure 2.**
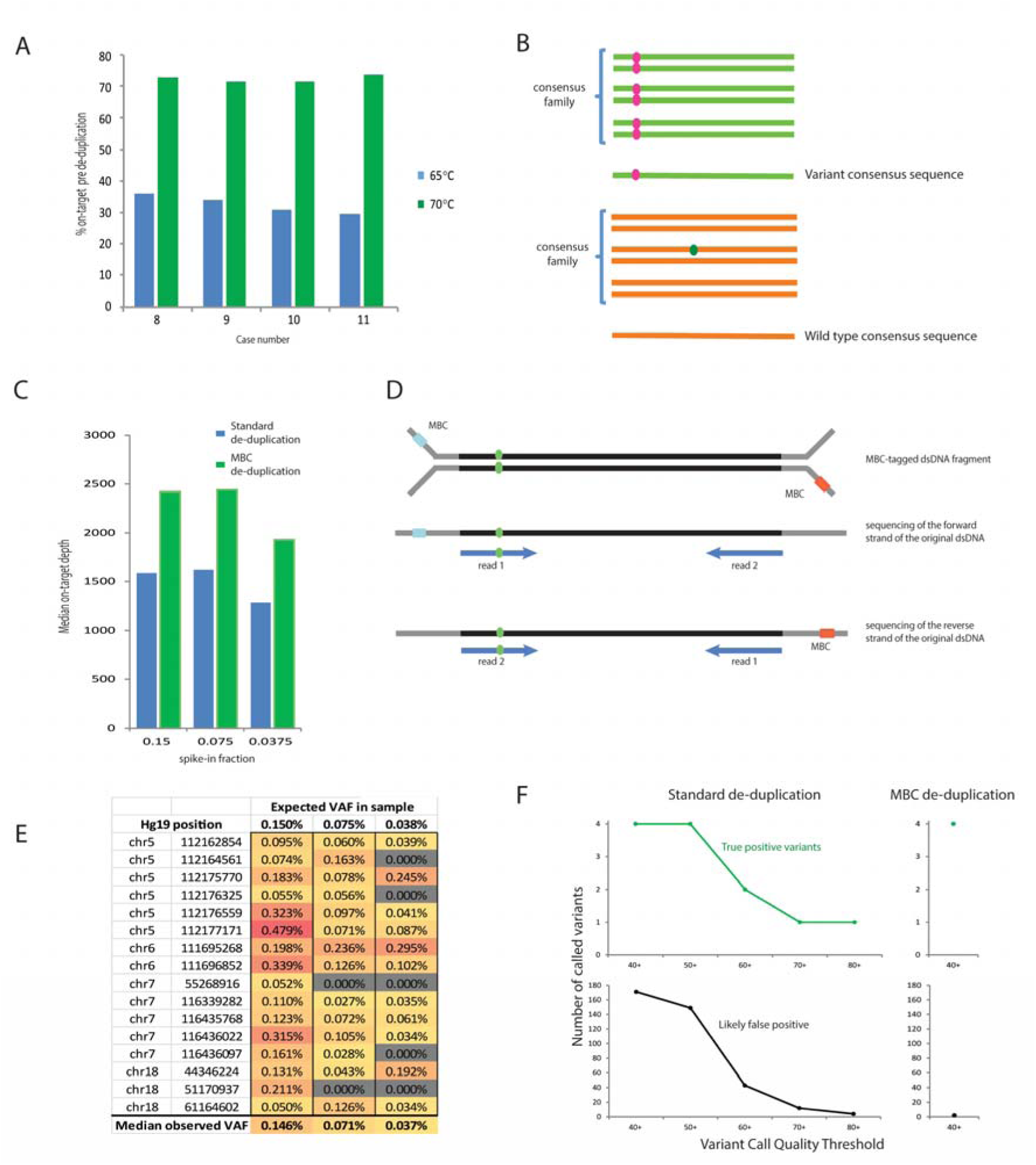
(A) Percentage of reads on-target before de-duplication in samples prepared with 65 C vs 70 C post-capture washes. (B) Graphic depicting the principles of MBC error correction. Reads with the same MBC that map to the identical genomic location are grouped into a consensus family. If a variant (pink) occurs in all reads then the consensus read sequence will be variant for that base (top). However if a variant (green) is only detected in a small fraction of the reads in the family, it will be disregarded and the consensus read sequence will be wild-type (bottom). (C) Median on target sequencing depth of the cfDNA mixing experiment after MBC de-duplication vs standard de-duplication. (D) Illustration of duplex read pair detection. A double stranded cfDNA fragment (black) containing a variant (green) is depicted, ligated to Y-shaped MBC-tagged adapters (grey). Each of the two strands of the cfDNA duplex molecule is labeled with a different MBC (blue or red boxes). The resulting paired-end sequencing data will be orientated differently for the two original strands. The original forward strand of the cfDNA duplex will result in read pairs where the forward read (relative to the reference genome) will be read 1 and the reverse read will be read 2. The reverse strand of the original cfDNA duplex will result in read pairs with the opposite orientation: i.e. the reverse read is read 1 and the forward read is read 2. Hence the order of the read pair sequences with respect to the reference genome alignment can be used to detect likely duplex DNA strands. (E) Expected and observed variant allele frequencies (VAF) and genomic positions for the 16 SNPs in the cfDNA mixing experiment. (F) Impact of MBC error correction on true positive and false positive calls. The top panels show the number of true positive variants (expected SNPs) that were bioinformatically called in the mixing experiment with standard de-duplication (left) and MBC de-duplication (right) using different variant call quality thresholds. The lower panel shows the number of likely false positive variant calls (not observed in the deep sequencing of either cfDNA sample used in the mix) for standard de-duplication (left) and MBC de-duplication (right).

We next used the MBCs to de-duplicate the sequencing data and to perform sequencing error correction. The SureCall software first groups sequencing reads that have identical MBCs and share the same genomic alignment position into a family (Figure 2B). For a family of reads that do not all have identical sequences, the software computes a score for each candidate sequence based on the mapping and base qualities in that read, which is then multiplied by the number of reads with identical sequences. The sequence with the highest resulting score is used as the consensus sequence in the merged read. This should reduce random errors arising during PCR and sequencing, as these are not usually common to all reads of a family.

After MBC de-duplication, the median on-target depth for the 70°C washed samples was 1,782x. This is theoretically sufficient to achieve a detection limit of approximately 1 mutated DNA fragment in 1,782 molecules (0.056%). However, the sensitivity for the de novo detection of mutations may be lower in practice, as more than one read is likely to be required to support robust bioinformatic variant calls. For example, if three independent reads were required, then the detection limit may be closer to 0.17% (3 reads in 1,782). Thus, we designed a mixing experiment to test the ability to reliably detect and also bioinformatically call mutations with such low frequency.

### Assay sensitivity and specificity

We used cfDNA from two donors that differed in 16 homozygous SNPs within our targeted region and prepared a dilution series with 0.15%, 0.075% and 0.0375% cfDNA from donor A spiked into the cfDNA from donor B. 25 ng of each mixture were then used for sequencing. A median of 74,030,118 reads were sequenced per sample, resulting in a median on-target depth of 21,651x before de-duplication. Sequencing data from each sample was then processed in two ways: First, we used MBCs for de-duplication and calling of consensus sequences. Alternatively, we performed standard de-duplication, which uses the genomic position of each read pair.

The median on target depth was higher after MBC de-duplication (2,420x with MBC de-duplication versus 1,587x with standard de-duplication; Figure 2C). This had been anticipated as DNA fragments in the cfDNA sample that map to the same genomic location will be tagged with distinct MBCs, as will the forward and the reverse strand of each double stranded ‘duplex’ DNA (Figure 2D). Standard de-duplication removes all but one of these types of reads as it cannot distinguish them from PCR duplicates. However, these reads are retained as independent units of information when MBCs are used.

We first investigated whether the spiked-in SNPs in this mixing experiment could be re-identified. All 16 were detected in the 0.15% mix, 14/16 (87.5%) were detected at 0.075% and 11/16 (68.75%) at 0.0375% mixing ratio (Figure 2E). The observed VAFs varied between the SNPs within each sample, most likely as a result of stochastic sampling at such ultra-low VAFs. Overall, this ultra-deep cfDNA sequencing assay allowed the robust detection of single nucleotide variants present at 0.15% and still showed a high sensitivity for the detection at 0.075% (i.e. one mutated DNA molecule out of 1,333 DNA molecules).

Our next aim was to assess whether MBC error correction improved the calling accuracy of ultra-low frequency variants, which is technically more challenging than the re-identification of known variants at specific positions. We applied the SureCall software to assess how MBCs influence the ability to detect any of the 9 SNPs from sample A that were present at a frequency of 0.15% in the cfDNA mixture (SNPs from sample B would not be informative as they were present in 99.85% of DNA molecules in the cfDNA mix and were hence always called accurately against the reference genome). Calling of the mixing experiment was first performed using MBC de-duplicated data and then the same data processed with standard de-duplication. The same call parameters were used for both. We also observed that true variants (i.e. the spiked-in SNPs) were always at least supported by two different consensus families that mapped to the same genomic position. These two families differed in whether the variant was seen in read 1 or read 2 during paired-end sequencing. These were highly likely to represent the two different strands of the original duplex cfDNA molecule, as when MBCs are ligated to the cfDNA fragment, the orientation of the MBC will be different for the two strands (Figure 2D). Based on this observation, we added the requirement that such a ‘duplex configuration’ has to be present in order to accept a mutation as genuine in our post call filtering steps. The presence of a variant in at least one more family with a different alignment position was also added to our post-call filters to assure high specificity. With these criteria, a variant had to be present in at least 3 consensus DNA families before it could be accepted as a mutation call in the MBC de-duplicated data. In order to enable a meaningful comparison, mutations in the standard de-duplicated data were also required to be present in at least three reads.

Mutation calling in the entire 163.3 kb target region of our assay, using standard de-duplication and a low stringency variant call quality threshold (VCQT) of 40 (manufacturer default setting: 100), detected 4 of the 9 spiked in SNPs (Figure 2F) but also generated 171 additional mutation calls. These additional variants are likely false positives as they were not identified by deep sequencing of the individual cfDNA samples. Stepwise increase of the VCQT first reduced the likely false positive calls to 149 while retaining all 4 true positive calls but a further reduction in false positives (to only 4 at the highest tested VCQT of 80) was accompanied by a loss of sensitivity with only a single spiked-in SNPs called.

When the same data were called using MBCs and a low stringency VCQT of 40 (Figure 2F), the same 4 spiked-in SNPs that had been found above were called but with only 2 additional and likely false positive variants. Thus, MBCs reduced the false positives compared to standard de-duplication from 149 to only 2 (reduction of 98.6%) while maintaining the maximum number of 4 called SNPs. We further assessed why the caller failed to pick up the 5 other SNPs. Each of these had a variant allele frequency below 0.1% (Figure 2E) and these cannot be called by SureCall as the lowest possible setting for the minimum VAF is 0.1%.

These results show that MBC error correction leads to a dramatic decrease of false positive mutation calls and enabled the identification of variants with a sensitivity of 0.15% with high precision.

### Concordance of cfDNA- and tumour-sequencing in 28 patients with metastatic CRC

We continued to sequence cfDNA from FOrMAT trial patients until 28 patients had been consecutively analysed. Seven out of these had been sequenced with our early protocol using 65°C wash temperature and 21 with our optimized protocol using 70°C. The median sequencing depth across samples was higher with the use of the more stringent conditions (1,205x on-target reads after MBC de-duplication with 65°C and 2,087x with 70°C).

For each case, we analysed the concordance and discordance of mutation calls within the target regions common to the sequencing assay applied to the tumour biopsy and our cfDNA sequencing panel. Tumour tissue of 23 cases had been sequenced with the FOrMAT NGS panel (Supplementary Table 1). The target regions of our cfDNA assay and the FOrMAT assay overlapped for all exons of 11 genes (*APC*, *TP53*, *KRAS*, *SMAD4*, *FBXW7*, *CTNNB1*, *TCF7L2*, *ATM*, *NRAS*, *MAP2K1*, *MAP2K2*) and for the specific exons harbouring mutational hotspots for 5 genes (*PIK3CA*, *PTEN*, *BRAF*, *EGFR*, *ERBB2*). Four tumour samples had been analysed with an amplicon sequencing panel used in routine clinical diagnostics that only included 5 genes (*BRAF*, *KRAS*, *NRAS*, *PIK3CA* and *TP53*), which were all present in our cfDNA sequencing panel. One case had failed any tissue sequencing and no mutations were known.

88% (80/91) of all mutations that had been found by tumour tissue sequencing were called in the cfDNA (Figure 3A). In 89% of cases (24 of 27) in which tissue and cfDNA had been analysed all previously known mutations were identified.

**Figure 3.**
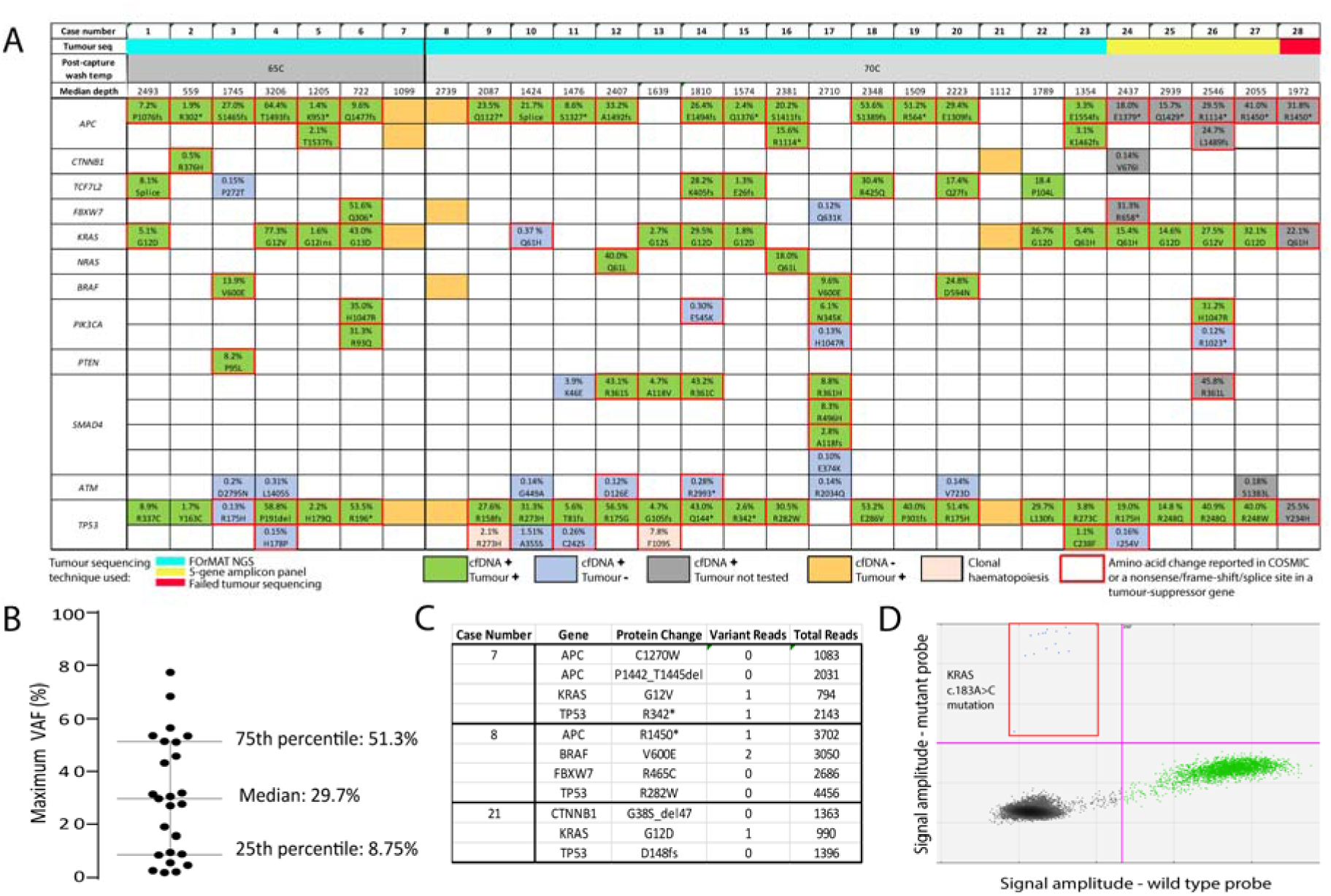
(A) Concordance of mutations identified by cfDNA sequencing and by sequencing of tumour material. Mutations identified in both cfDNA sequencing and tumour sequencing are coloured green. Novel variants called by cfDNA sequencing and not by tumour sequencing are coloured blue. Variants not detected by cfDNA sequencing that were detected in tumour sequencing are coloured orange. Pink indicates clonal haematopoiesis. Red outlines indicate mutations reported as tumorigenic in COSMIC. Variants in grey have been identified in the cfDNA of patients that either had been sequenced using the limited 5-gene amplicon panel or failed FOrMAT sequencing. Percentages indicate VAF in cfDNA. (B) Range of maximal VAFs across 28 cfDNA samples. (C) Read depth and number of consensus family reads supporting each of the 11 variants in cases 7, 8, and 21 that had not been called in cfDNA but had previously been detected in tumour tissue. (D) ddPCR validation of the *KRAS* c.183A>C mutation that results in the amino acid change Q61H in case 10. Green dots: droplets with wild-type DNA, blue dots (outlined by the red quadrant): droplets with mutant DNA, black dots: droplets that have no incorporated DNA.

All 11 mutations that were not called in cfDNA but had been identified in tumour tissue were from only 3 of the 28 cases (11%). Detailed analysis of the sequencing data of these cases showed that 5 out of the 11 mutations had been detected in only 1 or 2 reads each, equivalent to a median VAF of 0.066% which is below the detection limit of our assay (Figure 3C). Sufficient cfDNA was remaining from case 8 for assessment of mutation abundance with highly sensitive digital droplet PCR (ddPCR). Using ddPCR probes for the mutation encoding the BRAF V600E oncogenic variant, we identify 2,830 wild type DNA fragments but no mutated fragments (data not shown). Calculating the 95% confidence interval for a binomial distribution, this indicates that the true allele frequency is likely between 0% and 0.11%. This orthogonal validation confirmed a very low tumour-derived cfDNA fraction and hence a biological reason rather than technical failure explained the inability to detect this mutation.

We next assessed mutations called by cfDNA sequencing in genes that had not been sequenced in the corresponding tumour tissue. cfDNA sequencing detected *APC* mutations in each of the 4 cases (one in each of three cases and two in one case) whose tumours had only been analysed by the 5-gene amplicon panel. Furthermore, one mutation was found in each of *FBXW7*, *CTNNB1*, *ATM* and *SMAD4*. Our assay also detected mutations in *APC*, *TP53* and *KRAS* in case 28 that had previously failed tumour tissue sequencing attempts. Ten out of these 12 mutations (83%) encoded protein changes previously observed in cancer samples according to the COSMIC (21) cancer mutation database. Their recurrent occurrence in tumours suggests that these are likely driver mutations, further demonstrating that our assay is able to detect biologically important cancer mutations directly from cfDNA.

We subsequently analysed mutations called in cfDNA that had not been detected when the same gene had been analysed in tumour tissue. 23 such mutations were detected: 7 in *TP53*, 8 in *ATM*, 3 in *PIK3CA*, 2 in *SMAD4* and one each in *KRAS*, *FBXW7* and *TCF7L2*. 100% of mutations called in oncogenes (1 in *KRAS*, 3 in *PIK3CA*) were canonical activating cancer driver mutations. The other 20 mutations were located in tumour suppressor genes and 48% (8/20) of these were either nonsense mutations or encoded for amino acid changes that had previously been detected in cancer according to the COSMIC database, suggesting that these were also driver alterations. Overall, 56.5% (13/23) of mutations detected in cfDNA but absent in tumour were likely cancer driver mutations.

TP53 mutations in cfDNA do not necessarily originate from the tumour itself, but can be the result of a clonal expansion of blood cells (9), termed clonal haematopoiesis (22, 23). We assessed genomic DNA from blood cell pellets that had been sequenced with the FOrMAT panel, and found that 2 of the *TP53* mutations that had been detected in cfDNA but not in matched tumour tissue (Cases 9 and 13) were present in blood cells with VAFs of 2.1% and 7.8%, respectively (Supplementary Table 2). These are cases of clonal haematopoiesis, showing the importance of sequencing DNA extracted from blood to avoid interpreting such variants as cancer-associated mutations.

We next assessed the VAFs of the mutations that were called in cfDNA but not in matched tumour tissue. These were on average 105-fold lower than the VAF of the most abundant mutation detected in the same cfDNA sample (Supplementary Figure 1). The dramatically lower VAF measurements indicate that most of these mutations would have to be confined to small subclones in these tumours.

Similar to data showing that subclonal *APC*, *KRAS*, *NRAS* and *BRAF* mutations are extremely rare in untreated mCRCs (24), we only detected a single divergent alteration between cfDNA and tumour tissue sequencing among these genes in our series. The activating mutation in the *KRAS* gene (Q61H) was detected with a VAF of 0.37% in cfDNA of case 10 (Figure 3A) and none of 633 reads from the matched tumour showed the mutation. Assessing the clinical data of our cohort showed that this was the only case that had been treated with the anti-EGFR antibody cetuximab prior to blood collection. This subclonal *KRAS* mutation was hence a likely driver of acquired therapy resistance that had evolved during cetuximab therapy as previously described (25). ddPCR testing on leftover cfDNA from the same sample confirmed this mutation (Figure 3D), providing orthogonal validation that our technology is suitable for the detection of clinically relevant subclonal alterations.

In contrast to the high concordance observed for the above genes, driver mutations in PIK3CA were frequently discordant with 3 out of a total 7 mutations only detectable in cfDNA (E545K, H1046R, R1023*; Figure 3A). Two of these (cases 17 and 26) were likely cases of parallel evolution as other activating *PIK3CA* mutations had already been detected in those tumours and in cfDNA.

Finally, cfDNA sequencing also called mutations in the *ATM* tumour suppressor gene in 8/28 cases. Seven were found in cases where sequencing of DNA from matched tumour tissue showed wild type sequence only (data not shown) and one in the cfDNA of a case where the tumour had only been sequenced with the 5-gene panel that did not include *ATM*. All *ATM* mutations had very low VAFs (median: 0.17%) and only 2 out of the 8 variants encode protein changes catalogued in COSMIC. Notably, no *ATM* mutations were called in our reference set of 6 healthy donors, indicating that the mutation calls in cfDNA from mCRC patients are not the result of a generally high false positive call rate in this gene.

### Analysis of off target reads allow genome wide DNA copy number aberration analysis

Cancer genetic aberrations are not confined to point mutations but also include structural DNA aberrations and copy number changes. Thus, we investigated whether we could maximise the characterisation of each cancer in a single assay by also reconstructing genome-wide copy number aberration (CNA) profiles from our targeted cfDNA sequencing data. We applied the CNVkit algorithm (26) that can infer CNA profiles using off-target reads generated by targeted solution hybrid capture DNA sequencing and which are scattered across the genome. Genome-wide CNA profiles were derived for 20 of the 28 cases (71%) in our cohort. CNA profiling was successful from cfDNA samples where the maximum mutation VAFs were as low as 8.6% (examples in Figure 4A), showing that CNA profiling is possible in samples with intermediate to low tumour derived DNA contents. The 8 samples where only a flat CNA profile had been detected showed very low maximum mutation VAFs ≤5.6 %.

**Figure 4.**
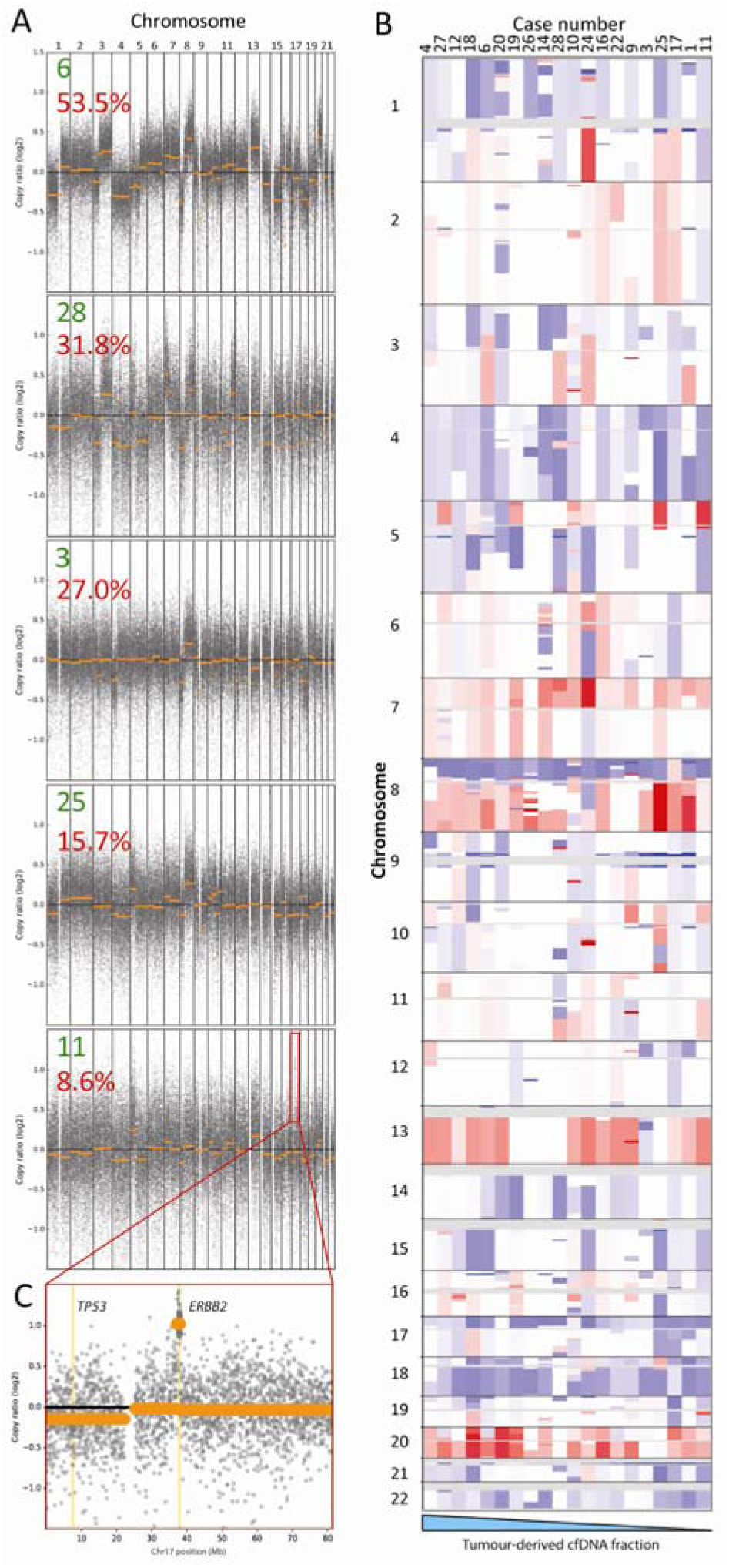
(A) Genome wide copy number aberrations can be detected from targeted cfDNA sequencing, even where tumour content is low. Representative log copy ratio plots for five cases (green number) in our cohort with tumour content ranging from 53.5% to 8.6% (red number indicates max VAF) are shown. (B) Genome wide heat map of segmented copy number raw log ratio data after amplitude normalization. Gains are red and losses are blue. Profiles are ordered (left to right) from highest to lowest tumour content (based on maximum VAF) for all 20 cases that had a visible CNA profile. (C) Focused log copy ratio plot of chromosome 17 for case 11 which had a high level amplification of *ERBB2*.

19 of 20 CNA profiles showed chromosome 17p loss and 18 out of 20 chromosome 18q loss (Figure 4B), which are the most frequent chromosome losses in CRC (27). Gains of chromosomes 1q, 7, 8q, 13 and 20, which are also common in CRC, were present in multiple samples. This shows that off-target reads from cfDNA sequencing can be used to reliably reconstruct genome wide DNA copy number profiles.

We also inspected high-level amplifications in these copy number profiles for potentially targetable alterations. A high level amplification involving the *ERBB2* oncogene was clearly visible in case 11 despite a low fraction of tumour-derived cfDNA in this sample (Figure 4C). The ERBB2 amplification had also been detected by the FOrMAT sequencing panel in the matched tumour, validating the ability to profile CNA with our cfDNA sequencing technology to detect actionable copy number aberrations. No other amplifications had been detected in the tumour biopsies with the FOrMAT NGS panel and no further targetable amplifications were apparent in the remaining cfDNA samples.

## Discussion

We described the development of an ultra-deep and error-corrected cfDNA sequencing protocol using off-the-shelf MBCs in combination with a custom-designed solution hybrid capture panel. A mixing experiment with cfDNA from two donors showed 100% sensitivity to detect variants with a VAF of 0.15% and that 87% of variants were still detected at a VAF of 0.075%.

The use of MBC error correction reduced false positive mutation calls by 98.6% while maintaining true positive calls. This accuracy was achieved by mutation calling followed by a set of post-call filtering steps that made use of the information originating from the MBC, as well as the background variant rates observed in cfDNA from a small number of 6 healthy donors. We therefore did not need complex error correction models requiring the sequencing of a large number of healthy donor samples, which are often impractical for many research applications, particularly those requiring frequently changing custom gene panels.

Our subsequent analysis of cfDNA from 28 patients with mCRC demonstrated that 88% of the cancer mutations that had been detected by clinical grade tumour tissue sequencing were also called in cfDNA. This sensitivity is similar to that reported for a single-gene cfDNA assay applied to CRC patients (87.2%) (1) and a 54 gene assay applied to multiple tumour types including CRC (85%) (15, 28).

Small insertions and deletions (indels) can be more difficult to call than point mutations, particularly when present at low abundance. Our cfDNA assay called 23 of 26 indels (88.5%) that were known based on tumour sequencing, showing a similar performance to point mutation detection (57 of 65 called; 87.7%). The called indels included 7 with VAFs <5% and a minimum VAF of 1.3%. The indel calling performance at lower VAFs could not as yet be tested due to a lack of cases with both known indels and a tumour derived cfDNA fraction below 1%.

cfDNA sequencing detected several additional cancer driver mutations that were not reported by tumour sequencing. Seven such variants were observed in TP53, all but one of which were recognized tumorigenic mutations listed in COSMIC. Two of them were also observed in the matched blood samples, indicating that these are the consequence of clonal haematopoiesis. The discovery of clonal haematopoiesis in 7% of our cohort demonstrates the need to sequence DNA from blood cells in parallel with cfDNA, in order to assess whether *TP53* mutations in cfDNA are likely to be of cancer origin.

In one patient, we detected a KRAS Q61H variant that was absent from the matched tumour. This patient had progressed on cetuximab therapy and treatment had been stopped 8 months prior to the cfDNA sample collection. It is therefore likely that this represents a drug resistant subclone that evolved during cetuximab therapy and remains detectable at low level.

Multiple *PIK3CA* activating mutations detected in two cases are likely to represent parallel evolution events. Parallel evolution strongly suggests that these alterations are selectively advantageous as shown for other tumour types (11). Importantly neither of these two patients had received prior anti-EGFR therapy thus these are not resistance driver mutations that evolved as a response to targeted therapy exposure as previously described (25).

Overall, mutations that were only called in cfDNA showed a high fraction of likely driver aberrations and a pattern that is consistent with previous mCRC evolution studies (24), such as the absence of divergent *RAS*/*RAF* mutations in cases not previously treated with anti-EGFR antibodies and frequent heterogeneity of *PIK3CA* mutations. This supports the notion that many of these are evolving driver mutations that could not be detected by the single time-point tumour biopsies. Further orthogonal validation, for example with ddPCR, could provide more detailed information on what fraction of these are true positives.

Together, these data show that our cfDNA analysis technology may address the subset of 20% or more of CRC patients who cannot be molecularly profiled due to unobtainable or inadequate biopsy tissues (17). Due to its minimally invasive nature, ultra-sensitive cfDNA sequencing can also be applied serially at multiple time-points, for example to monitor the evolution of subclonal drug resistance driver mutations in target gene panels without prior knowledge of specific loci where resistance mutations will occur. As the number of targeted therapies increases small custom target enrichment panels, which can be readily adapted for the tumour type and therapeutic agent in question, could be used to investigate the full tumour genomic landscape of point mutations, indels and CNAs. This would facilitate both the identification of novel resistance mechanisms and help to further characterise cancer evolution.

In conclusion, this cfDNA sequencing approach with customizable and off-the-shelf reagents showed a similar sensitivity and specificity as published cfDNA sequencing techniques using bespoke reagents and more complex analyses. Performance from small cfDNA quantities that can be obtained from most patients with mCRC particularly enable longitudinal studies for monitoring and detection of evolving mutations.

## Methods

### Patients and samples

Samples from 6 healthy donors (HD) were collected after obtaining written informed consent through the Improving Outcomes in Cancer biobanking protocol at the Barts Cancer Centre (PI: Powles), which has been approved by the UK national ethics committee (approval number: 13/EM/0327). 27 ml of blood were collected from each donor during a single blood draw.

Blood samples from 58 patients with metastatic colorectal cancer were acquired after obtaining written informed consent through the FOrMAT clinical trial (Feasibility Of Molecular characterization Approach to Treatment, PI: Starling, ClinicalTrials.gov NCT02112357) at the Royal Marsden Hospital, which has been approved by the UK national ethics committee (approval number: 13/LO/1274RM). FOrMAT is a single centre translational study assessing the feasibility of next generation sequencing to guide treatment personalisation in patients with advanced gastrointestinal tumours.

Archival or fresh tumour specimens from patients enrolled in the FOrMAT trial had been sequenced with solution hybrid capture enrichment of 46 cancer driver genes relevant for gastrointestinal cancers as described (17). Mutation calls, BAM files of tumour and blood sequencing data as well as clinical data were available for our analyses.

### Blood sample processing and cfDNA extraction

Blood was collected in EDTA blood tubes and centrifuged within 2 hours for 10 minutes at 1600g. The plasma layer was carefully removed and stored at −80°C. Upon thawing, samples were centrifuged for 10 minutes at 16000g and 4°C to remove debris. cfDNA was then extracted from 4 ml plasma per mCRC patient and from 2×4 ml from healthy donors using the Qiagen QIAamp Circulating Nucleic Acid Kit, following the manufacturer’s protocol. cfDNA was eluted in 30 μl 10mM Tris Ultrapure buffer + 0.1 mM EDTA (low EDTA TE) and stored at −20°C until library preparation. cfDNA was quantified on a Bioanalyzer High Sensitivity DNA chip, and if this showed a high concentration, on a 7500 DNA chip (Agilent).

### cfDNA sequencing

We modified the Agilent SureSelect XT HS protocol (details below) in order to assure a reliable performance with 25 ng cfDNA input. All PCR steps were performed on an Eppendorf Mastercycler nexus GSX1. 8 cycles of pre-hybridization PCR were optimal for 25 ng of input cfDNA to generate the amount of product required (500–1000ng) for the in-solution capture step. The entire product was used as input for hybridization to the custom-designed Agilent SureSelect capture bait library, targeting 32 genes as well as 40 SNP positions on chromosome 18q (target region: 163.3kb). All other reagents were added according to the manufacturer's protocol and 60 cycles Fast Hybridization were performed. Capture was started immediately after the final hybridization cycle and proceeded for 30 minutes at room temperature.

Post-capture washes were performed with two different conditions. The manufacturer’s initial protocol recommended 3 incubations of 10 minutes each at 65^o^C. We subsequently used more stringent post-capture washes with 6 incubations of 5 minutes each at 70^o^C. This optimized cfDNA library preparation protocol was used for the remaining 23 mCRC patients and 6 further healthy donors.

10 PCR cycles were used for Post-Capture Amplification and this was followed by two rounds of 1x Ampure XP bead cleanup to remove un-incorporated primers. The final prepared sequencing libraries were checked on a Bioanalyzer High Sensitivity DNA chip and quantified by qPCR using the Kapa Library Quantification kit before pooling. Pools were clustered using an Illumina cBot and sequenced with paired end 75 reads on an Illumina HiSeq2500 in Rapid output mode.

### Variant calling

Agilent SureCall (version 4.0.1.45) was used to trim and align fastq reads to the hg19 reference genome with default parameters. Raw on-target depths and the on target fraction were assessed before de-duplication. MBC de-duplication was performed with SureCall permitting one base mismatch within each MBC and consensus reads that were only supported by a single read were removed as MBC error correction is ineffective on such ‘lone reads’. The on-target depth after de-duplication was assessed after single read families were removed. Variant calling was performed using the SureCall SNPPET SNP Caller with the following parameters: Variant score threshold = 0.01; Minimum quality for base = 30; Variant call quality threshold = 40; Minimum Allele Frequency = 0.001; Minimum number of reads per barcode = 2; no region padding; and masked overlap between reads. Further filtering of the primary calls was performed after visualizing sequencing data on the Integrated Genome Viewer (IGV)(29). Variants only passed filtering if supported by a) at least one duplex pair of consensus families in which the variant was not the last base in the read and b) at least two reads with different alignment positions. Variants predominantly located in reads with an alignment score of zero or in reads with multiple non-contiguous non-reference bases were removed as these are usually indicative of misalignment. Standard de-duplication and subsequent calling was also performed with SureCall using the same parameters, except that de-duplication was solely based on genome alignment and insert size. As the post call filtering of MBC de-duplicated data essentially required a variant to be present in a minimum of three reads (two that form a duplex and one additional read with a different alignment position), we also mandated a minimum of three reads with a specific variant to support a mutation call in the standard-duplicated data to enable a fair comparison. All variant positions identified in patient cfDNA were checked in the dataset of six HD samples using bam-readcount (https://github.com/genome/bam-readcount) in order to filter out any false positives, such as due to misalignment. The majority of called variants were completely absent in the six HD samples (Supplementary Table 3), however, mutations whose VAF was not at least double that of an identical variant in a healthy donor sample were discounted.

### Genome-Wide Copy Number Aberration Analysis

Bam files resulting from MBC de-duplication before removal of single-read consensus families were used to generate genome-wide DNA copy number profiles using CNVkit (26). The six sequenced healthy donor cfDNA samples were used as a pooled normal reference dataset. Antitarget average size was set to 30,000 bp to improve resolution. Seg files were created from the .cns segmented output for visualisation of the detected copy number aberrations as a heatmap on IGV.

### ddPCR

cfDNA from 2 patients was probed for *BRAF* or *KRAS* variants identified from tumour sequencing, using commercially available ddPCR SNP Genotyping Assays (Life Technologies; Assay ID *A44177 BRAF_476* and A44177 *KRAS*_555). Input cfDNA amount varied, depending on residual material available (20 ng for case 8 and 17 ng for case 10). cfDNA was added to a ddPCR reaction containing 11 μl mastermix (10 μl 2x ddPCR Supermix for Probes (no dUTP) and 1 μl 20x target primer/probe mix for both mutant and wild type alleles) and made up to a total volume of 20 μl with nuclease-free water.

The reaction was partitioned into a median of 17,000 droplets per sample in a Bio-Rad QX-200 droplet generator according to the manufacturer’s protocol. Emulsified PCR reactions were run on a 96 well plate on a G-Storm GS4 thermal cycler incubating the plates at 95°C for 10 minutes followed by 40 cycles of 95°C for 15 sec and 60°C for 1 minute. Plates were read on the QX200 droplet reader using QuantaSoft analysis software (Bio-Rad) to acquire and analyse data. At least four positive and negative control wells were included in every run to verify assay performance and facilitate thresholding in fluorescence values. For each patient, plasma was analysed in triplicate and ddPCR results are based on the combined data of these wells.

## Acknowledgements

We would like to thank all patients participating in the FOrMAT clinical trial and the clinical research team members at the Royal Marsden Hospital who supported the sample collection. The study was supported by charitable donations from Tim Morgan to the Institute of Cancer Research, from Philip Moodie to The Royal Marsden Cancer Charity and by a Clive and Ann Smith Fellowship. The study received funding by CRUK, a Wellcome Trust Strategic Grant (105104/Z/14/Z), the Royal Marsden Hospital/Institute of Cancer Research National Institute for Health Research Biomedical Research Centre for Cancer and by a CRUK Clinical PhD Studentship.

